# Chromatin enables precise and scalable gene regulation with factors of limited specificity

**DOI:** 10.1101/2024.06.13.598840

**Authors:** Mindy Liu Perkins, Justin Crocker, Gašper Tkačik

## Abstract

Biophysical constraints limit the specificity with which transcription factors (TFs) can target regulatory DNA. While individual nontarget binding events may be low affinity, the sheer number of such interactions could present a challenge for gene regulation by degrading its precision or possibly leading to an erroneous induction state. Chromatin can prevent nontarget binding by rendering DNA physically inaccessible to TFs, at the cost of energy-consuming remodeling orchestrated by pioneer factors (PFs). Under what conditions and by how much can chromatin reduce regulatory errors on a global scale? We use a theoretical approach to compare two scenarios for gene regulation: one that relies on TF binding to free DNA alone, and one that uses a combination of TFs and chromatin-regulating PFs to achieve desired gene expression patterns. We find, first, that chromatin effectively silences groups of genes that should be simultaneously OFF, thereby allowing more accurate graded control of expression for the remaining ON genes. Second, chromatin buffers the deleterious consequences of nontarget binding as the number of OFF genes grows, permitting a substantial expansion in regulatory complexity. Third, chromatin-based regulation productively co-opts nontarget TF binding for ON genes in order to establish a “leaky” baseline expression level, which targeted activator or repressor binding subsequently up- or down-modulates. Thus, on a global scale, using chromatin simultaneously alleviates pressure for high specificity of regulatory interactions and enables an increase in genome size with minimal impact on global expression error.

**Significance Statement:** Reliably keeping a gene off is as important as controlling its expression level when the gene is on. Yet both tasks become challenging in the packed nuclear environment of a eukaryotic cell, where the numerous and diverse regulatory proteins that are present cannot bind enhancers for target genes with perfect specificity. While regulatory schemes based on prokaryotic models would be overwhelmed by errors in such conditions, we show that chromatin-based regulation, an evolutionary innovation of eukaryotic cells, successfully rescues precise gene expression control by reliably keeping desired genes off. Our systems-level computational analysis demonstrates that this result is nontrivial, because chromatin opening must itself be correctly regulated. We furthermore identify when and how chromatin-based regulation outperforms alternative schemes.

---

Many biological functions require certain subsets of genes to be active independently of others. The targeted activation or repression of gene expression is achieved by regulatory factors including transcription factors, which must specifically locate and bind the regulatory regions for their target genes. However, some level of nontarget binding is unavoidable due to the chemical nature of protein-DNA interactions (1–5). Organisms have therefore evolved several different mechanisms for increasing the ratio of target to nontarget binding. In prokaryotes, transcription factors bind almost uniquely to their cognate binding sites, which are long enough to appear only very rarely by chance in the genome. Paradoxically, though the eukaryotic genome is two to three orders of magnitude larger than the prokaryotic genome, binding sites in eukaryotes tend to be shorter and more degenerate than in prokaryotes and, consequently, cannot specify unique genomic locations (6). This biophysical observation has motivated researchers to propose alternative means for eukaryotes to maintain targeted gene regulation, including cooperativity (7–9), clustering of low-affinity sites (6, 10–13), kinetic proofreading (14–18), and combinatorial control of transcription (9, 19–22).

One notable possibility for reducing unwanted gene expression involves the use of chromatin, the physical packaging of DNA by nucleosomes and associated proteins largely unique to eukaryotes (23). Nucleosomes may physically obstruct transcription factor binding to DNA wrapped around the core (24–26) or directly inhibit transcription initiation by blocking the assembly of the preinitiation complex (27–31). Longer-term silencing of gene expression additionally requires specific patterns of DNA methylation (32), histone modifications (33), and/or higher-order chromatin structures to maintain a transcriptionally repressive state (34–36). Although condensed or compacted chromatin may hinder large regulatory factors or complexes from accessing binding sites (26, 37–40) and most transcription factors, if they manage to bind nucleosomal DNA, will dissociate quickly (26), some factors are capable of reversing chromatin-based repression by recruiting other factors to decompactify chromatin (41–43) or evict, displace, or disassemble nucleosomes from DNA (38, 44). Such transcription factors with “pioneering activity” (PFs) induce the dynamic changes to genetic silencing that are associated with proper cell fate specification during embryonic development (32, 34, 45, 46). Whether “pioneer factors” are an entirely distinct category from “ordinary” transcription factors (47), or whether TFs simply vary in their capacity to induce “pioneering activity” based on context, remains a subject of open discussion (48).

Introducing chromatin alleviates some nontarget transcription factor binding by rendering DNA inaccessible— but not without a cost. In order to selectively (de)silence genes, chromatin remodelers and PFs must themselves be able to target certain binding sites in chromatin while ignoring others. Indeed, it has been proposed that PFs may enhance the recruitment of chromatin remodelers to appropriate nucleosomes (16) as part of an energy-consuming kinetic proofreading scheme (49, 50), variants of which may improve specificity by a factor of 300-400 over equilibrium schemes (51). The relationship between target specificity and chromatin appears even more intricate in light of recent work indicating that chromatin can improve the functional specificity of proteins with otherwise degenerate DNA binding domains, for instance the Hox class of homeodomain proteins (7, 52). Thus, chromatin-based transcriptional regulation may contribute in nontrivial ways to the resolution of the binding site specificity paradox in eukaryotes (12).

While many works have considered chromatin-based transcriptional regulation at individual loci (24, 29, 42, 43, 53–59), we are unaware of any that consider the *global* implications of using a combination of chromatin and free DNA-binding transcription factors for gene regulatory programs. The omission is notable for two reasons. First, using both chromatin and transcription factors enables new modes of genetic regulation spanning multiple genes or regulatory regions; for example, the same TF could be reused in multiple networks simply by changing the accessibility of its target regulatory regions. Second, limited binding site specificity constitutes a challenge only insomuch as multiple factors and multiple nontarget sites are present simultaneously; in other words, it emerges as a nontrivial constraint only when considered at the global level (9).

Here, we address this knowledge gap by employing a theoretical approach. Our focus is on global gene expression patterning: the task of accurately modulating individual expression levels for multiple active (ON) genes simultaneously, while keeping subsets of silent (OFF) genes from expressing altogether. We ask whether gene expression patterning benefits from chromatin-based mechanisms relative to regulatory schemes relying on TFs alone, and find biophysically realistic regimes of operation where chromatin lowers gene expression patterning errors by one to two orders of magnitude. The systems-level benefits we identify in this work synergize with the recently identified evolutionary benefits of chromatin-based regulation, suggesting how such complex schemes could have evolved and become selectively maintained (60).

## Model setup

Our model of chromatin landscaping and transcriptional regulation is deliberately simple, retaining only the key features of gene expression patterning that can be treated independently of the exact molecular implementation. Where possible, we have chosen parameter values to match measurements or estimates available in the literature.

We first define a gene regulatory architecture and then comparatively evaluate the ability of that given architecture to realize target patterns of gene expression in two scenarios: *free DNA* and *chromatin*. Specifically, for simulated genes of interest we will compute how precisely they could be driven towards different target expression patterns, assuming that the concentrations of their regulating factors could be optimized without constraint towards each target (Figure 1). Because no practical control scenario could ever outperform the mathematical optimum, the residual (minimal) *global expression error* (GEE) of this “best-case” outcome is a well-defined and theoretically justified performance measure – a loss function – for each scenario, introduced in previous work as “regulatory crosstalk” (9).

**Fig. 1.**
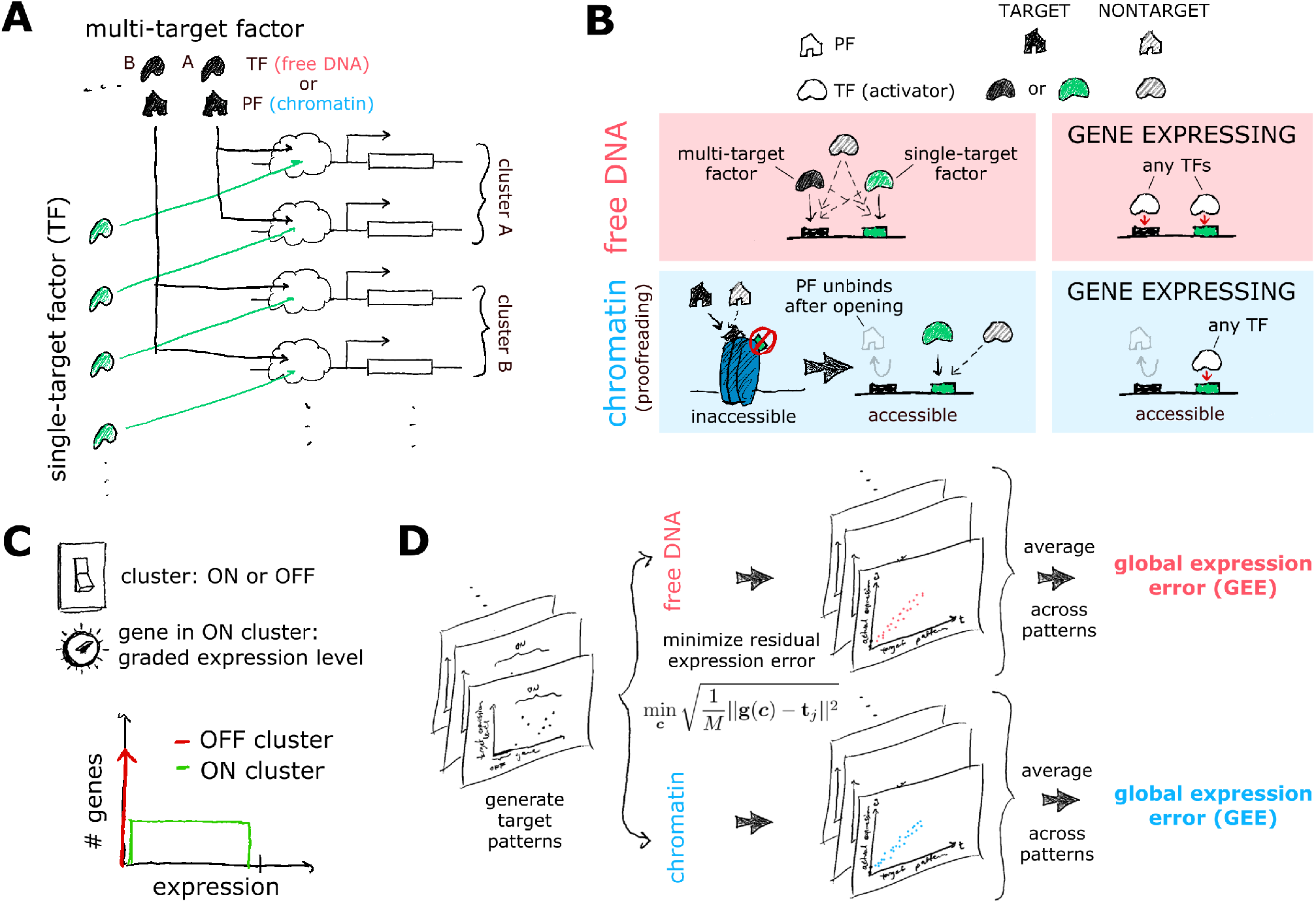
Model schematic. (**A**) Each gene is targeted for regulation by one multi-target factor (black; house-shaped PF in chromatin, bean-shaped TF in free DNA) and one single-target factor (green; TF). Genes targeted by the same multi-target factor are said to belong to the same cluster. (**B**) PFs (house shapes) and TFs (bean shapes) bind target and nontarget sites on inaccessible chromatin and free DNA/accessible chromatin, respectively, leading to chromatin opening or gene expression. (**C**) A target pattern is randomly generated by assigning all genes in the same cluster to the same ON or OFF state. OFF genes have zero target expression (red), while ON genes have randomly drawn graded expression, here a uniform distribution (green). (**D**) Global gene expression error (GEE) measures the patterning accuracy of a regulatory scenario (*free DNA* in red vs *chromatin* in blue) given a fixed architecture (which gene is regulated by which factors, as in panel A). GEE is calculated by optimizing regulatory factor concentrations to achieve target expression patterns as closely as possible for all genes simultaneously. Expression levels are calculated based on mathematical models for TF and PF binding (see Section S1).

Gene regulatory architecture consists of specifying how genes are targeted and regulated by various factors. Experimental evidence indicates that clusters of genes share temporal patterns of chromatin accessibility (61) and TFs outnumber factors with strong pioneering activity (62, 63). We therefore consider two classes of regulatory factors: *single-target* and *multi-target*. A single-target factor targets a single gene (i.e., there is a one-to-one mapping from single-target factors to genes), while a multi-target factor targets multiple genes that together form a co-regulated cluster. In our model, clusters are fixed and identical in size and each gene is targeted by exactly one single-target and one multi-target factor (Figure 1A). We initially assume all regulatory factors behave as activators and lift this assumption later.

We compare two regulatory scenarios: (1) the *free DNA scenario*, in which both the single-target and the multi-target factors are TFs; and (2) the *chromatin scenario*, in which the single-target factors are TFs and the multi-target factors are PFs. In the free DNA case, both TFs need to be bound simultaneously to activate a gene, but there is no sense of temporal ordering—that is, it does not matter whether the multi-target or single-target factor binds first. The expression level of a gene is thus simply the probability that both the single-target and multi-target factors are bound. In contrast, in the chromatin scenario, PFs must bind first to open enhancer chromatin (35, 61, 64) before TFs can bind to modulate transcription levels (Figure 1B); gene expression is thus the product of the steady-state probabilities that chromatin is permissive and the single-target TF is bound (35, 61, 64). This scenario thus does not require simultaneous binding by PFs and TFs at a regulatory locus. An identical genetic architecture can then be assessed comparatively in both the free DNA and chromatin scenarios, simply by changing the model for multi-target factor binding from TFs to PFs, respectively (Section S1).

In principle, transcription or chromatin opening can be initiated by any binding event—be it “target” or “nontarget”— albeit with different probability. To account for this observation, we assume that nontarget binding is weaker than target binding and occurs only among sites of the same type; that is, TFs cannot bind PF sites and vice versa. In contrast to the chromatin scenario, the free DNA scenario therefore experiences nontarget binding between single- and multi -target factors. The two scenarios also differ in their ability to distinguish between target and nontarget binding. While TFs follow a typical thermodynamic (equilibrium) binding scheme, we adopt a kinetic proofreading scheme for chromatin opening that allows for stronger rejection of nontarget binding by PFs than is possible at equilibrium (Figure S1). Kinetic proofreading expends energy to pass through at least one intermediate transition step between initial PF binding and final transcriptionally permissive state, e.g., by recruiting remodelers to stabilize chromatin in an open conformation (5, 65). To enable fair comparison between the two scenarios, we assign to all factors, TFs and PFs alike, the same intrinsic dissociation constants: *K*_*T*_ for their target and *K*_*NT*_ for all other (nontarget) sites. The ratio *S* = *K*_*NT*_ */K*_*T*_ is the *intrinsic specificity* of the factors; we vary it between simulations by changing *K*_*NT*_ . Our model is agnostic to whether nontarget binding arises from low-affinity binding, from truly nonspecific TF-DNA interactions, or by some other molecular mechanism. Further details of the models may be found in Section S1.

We can now mathematically define the *global expression error* (GEE), our main metric for comparing the “expressive capacity” of different regulatory scenarios. To compute the GEE, we begin by generating *N*_*T*_ random target gene expression patterns **t** = {*t*_*i*_} for genes *i* = 1, …, *M* according to two rules: (1) all genes in the same cluster (i.e., sharing a multi-target factor) are randomly chosen to either not express (OFF, *t*_*i*_ = 0) or express (ON, *t*_*i*_ > 0), such that the number of ON clusters is held fixed; (2) each gene within an ON cluster is independently assigned a random non-negative expression level from a uniform distribution. Thus, target expression levels across all genes follow a “spike and slab” distribution (Figure 1C), qualitatively recapitulating observations in early embryos (66). For each scenario (free DNA or chromatin) and each target pattern, the concentrations **c** of regulatory factors (PFs and TFs) are optimized to reduce the root mean square error between target patterns ***t*** and expression levels ***g*** realized by our model. The residual RMS error averaged over many target patterns finally yields the *global expression error* (Figure 1D):

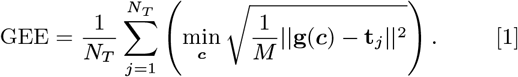

Unless stated otherwise, we report results averaged over sets of *N*_*T*_ = 100 target patterns that are shared between the two scenarios for fixed intrinsic specificity *S* = 1000, with the number of genes *M* fixed at 250. Multi-target factors each regulate 10 genes, and the number of ON clusters is fixed at 8 (80 active genes). We commonly analyze 5-fold variation around *S* = 1000 to visualize trends. Remaining parameters can be found in Table S1.

## Results

### Chromatin outperforms free DNA in expressive capacity

We optimized the concentrations of all regulatory factors for each target gene expression pattern to minimize GEE (Equation 1), both in the chromatin as well as the free DNA scenario. At high intrinsic specificity of the factors (*S* ≫ 10^3^), target patterns are achieved with high accuracy in both scenarios. Yet as *S* decreases, the free DNA scenario suffers visibly larger deviations from target expression levels compared to chromatin (Figure 2A). This is reflected in the respective GEEs of the two scenarios (Figure 2B), where chromatin outperforms free DNA for every *S*, with highest (10 − 50-fold) reductions in error seen at realistic values of *S* ≳ 10^3^. This effect is so stark that the intrinsic specificity of regulatory factors in the free DNA scenario would have to be ∼ 5× as high compared to the chromatin scenario to become competitive.

**Fig. 2.**
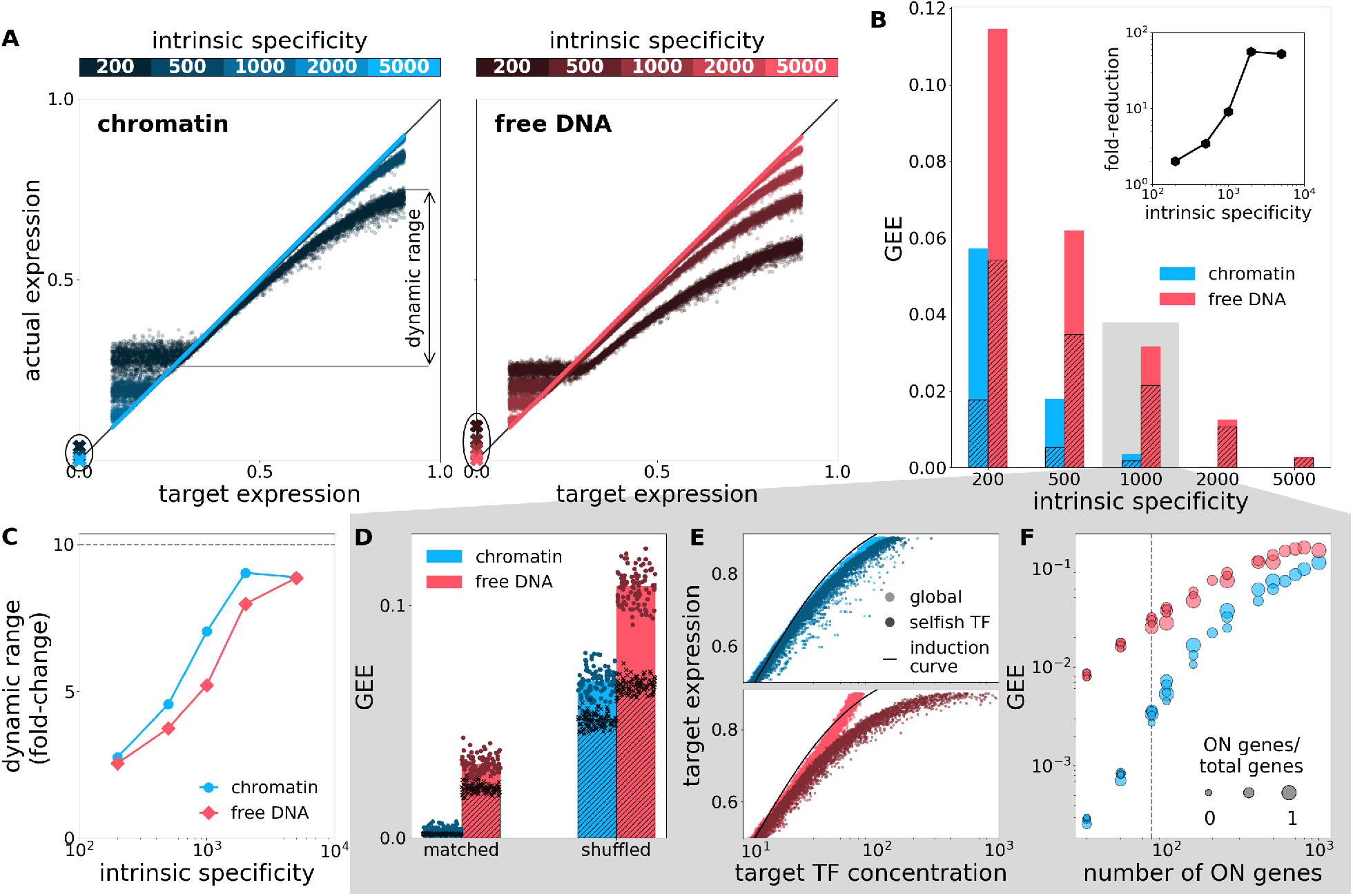
Chromatin achieves smaller global expression error compared to free DNA. (**A**) Actual vs. target expression levels after optimization of regulatory factor concentrations for chromatin (blue, left) and free DNA (red, right) for different intrinsic specificities *S* (colorbar, top). Points are ON genes in individual target patterns; crosses are OFF genes. Compared to free DNA, cromatin achieves expression levels for OFF genes much closer to zero (circled). (**B**) Global expression error is lower for chromatin (blue) than free DNA (red). Hatching indicates the GEE fraction due to erroneous expression of OFF genes. Inset: GEE fold-reduction for chromatin vs. free DNA. (**C**) Dynamic range (denoted in A) is greater for chromatin than free DNA. Dashed line indicates the maximal target dynamic range for this simulation. (**D**) GEE is lower for regulatory architectures where genes in the same cluster are either ON or OFF together (“matched”; left) compared to when OFF genes are randomly assigned to clusters (“shuffled”; right). Points are individual target patterns; bars are as in B. (**E**) Chromatin decouples regulatory control over ON genes. Light dots are individual gene expression levels achieved via global optimization over all PF and TF concentrations. Dark dots are optimization over individual single-target TF concentrations (see text). Ideal single-gene induction curve shown in black. (**F**) GEE is primarily determined by the number of ON genes rather than the absolute genome size or fraction of active genes (circle sizes). Dashed line indicates number of ON genes used for analyses in the other subpanels.

Detailed analysis reveals that two effects predominantly contribute to the substantial improvements in expressive capacity of chromatin over free DNA. First, chromatin much more effectively silences the genes that are supposed to be OFF, i.e., their expression can be regulated closer towards the 0 target (Figure 2A, circled crosses); this is reflected in a lower contribution to GEE from the OFF genes (Figure 2B, hatched) compared to free DNA. Second, while regulators in both free DNA and chromatin have difficulty inducing genes towards high expression levels, the failure at limited intrinsic specificity is much more pronounced in the free DNA scenario. This is reflected in its significantly smaller dynamic range of regulation—the mean difference between the highest and lowest achievable expression for the ON genes—compared to chromatin (Figure 2C).

Intuitively, the benefits of chromatin silencing should be maximized when the regulatory architecture groups genes into clusters that turn ON or OFF together (“matched” architecture), such that each cluster is regulated by a dedicated PF and TFs are employed solely to fine-tune the precise expression levels of ON genes. Our analysis in Figure 2D confirms this intuition: in the “matched” architecture, the GEE is 3-fold lower already for the free DNA scenario compared to its “shuffled” control, with the difference growing to more than 10-fold for chromatin. The shuffled architecture can no longer protect OFF genes: the need to express ON genes forces PFs to open chromatin, thereby exposing OFF genes in mixed clusters to regulatory influences from nontarget binding, which increases GEE. Therefore, for optimal expressive capacity, the genetic architecture needs to be matched to the co-activation structure of the desired target expression patterns.

Chromatin-based regulation has another important and biologically relevant benefit: it decouples and modularizes the regulation of ON genes. Consider the following. At global optimum, no gene will have perfectly achieved its target expression level. Suppose we choose a “selfish” gene that, starting from this global optimum, must reach its target expression level at any cost to other genes. The simplest way to regulate the selfish gene is to modulate the concentration of its corresponding single-target TF. Figure 2E shows that, in the chromatin case, we need not change the single-target TF concentration very much to appease the selfish gene: the required change is in line with the effective optimal induction curve of individual genes. In contrast, in the free DNA scenario, the concentration of single-target TF must be significantly increased if the “selfish” gene is to reach a high target expression level, at a cost of increasing error due to nontarget binding elsewhere. Chromatin-based regulation mitigates this effect by leveraging a division of labor between PFs and TFs, where PFs keep OFF genes OFF nearly without fail, while each TF modularly changes the expression level of its target ON gene without much impact on other expression levels.

Owing to this division of labor between PFs and TFs, chromatin retains an absolute advantage in GEE across a broad range of regulatory network sizes. The error tends to track the number of ON genes and varies little with the number of OFF genes (Figure 2F). For a fixed error, chromatin admits a much larger number of ON genes than free DNA, across two orders of magnitude in absolute genome size. Thus, regulating gene expression through chromatin accessibility could conceivably support the evolution of larger genomes with more genes active simultaneously, even if the intrinsic specificity of binding factors remains fixed.

### Optimal chromatin-based regulation co-opts nontarget binding

In all globally optimal solutions, genes with low target expression levels tend to be overexpressed relative to their targets, except at the highest specificities *S* that we consider (Figure 2A). This excess expression is almost entirely due to the high prevalence of nontarget binding (Figure 3A): already for ON genes with a middling desired expression level of 0.5, nontarget binding of single- and/or multi-target factors drives about 14% of expression in chromatin and 20% in the free-DNA scenario (Figure 3B). This observation led us to wonder whether globally optimal solutions that minimize GEE also implicitly minimize nontarget binding and vice versa, or whether GEE minimization perhaps productively co-opts nontarget binding to achieve target expression levels.

**Fig. 3.**
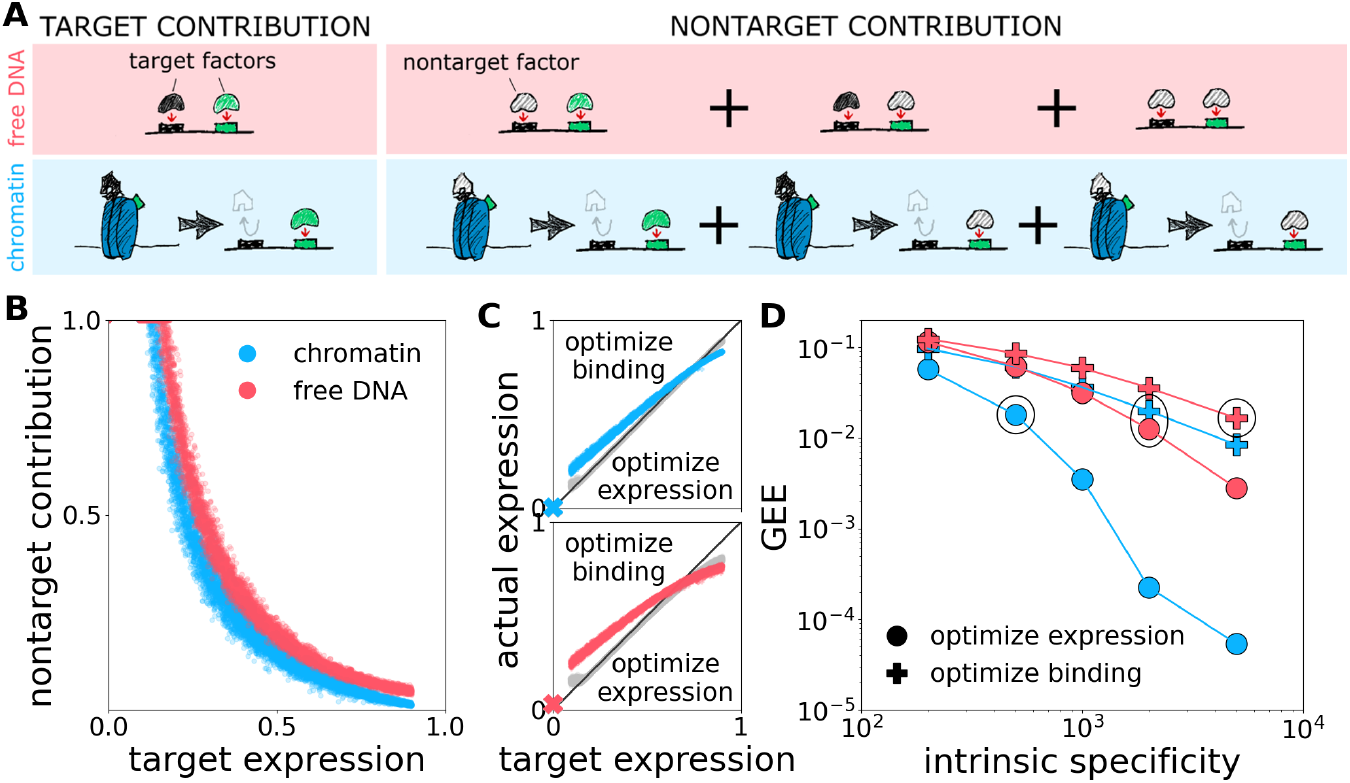
Co-opting nontarget binding contributes substantially to gene expressive capacity. (**A**) Nontarget binding contributions are defined as instances of gene expression in which the binding of at least one factor is nontarget. (**B**) Fraction of expression level (vertical axis) attributable to nontarget vs. target binding for chromatin (blue) and free DNA (red). Dots are individual genes pooled across target patterns with *S* = 1000. (**C**) Expression levels attained by optimizing to reduce global expression error (as in Figure 2, gray) compared to optimizing for target binding and against nontarget binding (color). (**D**) GEE is minimized when intrinsic specificity is high, chromatin is present, and nontarget binding is co-opted (blue circles). Nevertheless, multiple combinations of intrinsic specificity and regulatory scheme will often suffice to yield a very similar GEE (e.g., the circled points, see text).

To test the co-option hypothesis, we again turned to optimization. However, instead of optimizing factor concentrations towards matching target expression levels by any kind of binding, target or nontarget, as before, the new analysis only counts target binding as correct and all nontarget binding as erroneous regulation (see Section S2.1). We find that this alternative strategy based on target binding alone does allow graded expression of ON genes, but systematically over- (under-)expresses genes with low (high) target levels (Figure 3C). As a result, the GEE is substantially increased relative to the original scheme across the entire range of intrinsic specificities *S*; this effect is particularly pronounced in the chromatin scenario, where penalizing nontarget binding can increase GEE by up to two orders of magnitude (Figure 3D). Thus, optimizing for globally accurate gene expression patterning *is not* synonymous with maximizing regulation by target factors and minimizing the influence of nontarget factors. Rather, when intrinsic specificity is limited, co-opting nontarget binding is the most effective way to minimize global gene expression error.

Multiple regulatory strategies can achieve comparable expressive capacity. For example, a chromatin scheme that co-opts nontarget binding at intrinstic specificity *S* = 500 has a similar GEE as (1) a free-DNA scheme with *S* ≈ 2000 that co-opts nontarget binding; (2) a chromatin scheme with *S* ≈ 2000 that penalizes nontarget binding; and (3) a free-DNA scheme with *S* ∼ 5000 that penalizes nontarget binding (Figure 3D). This observation suggests a route by which chromatin might have alleviated the evolutionary pressure to maintain high intrinsic specificity: Starting from the free DNA scenario, if chromatin emerged due to selective forces not directly related to gene regulation (e.g., to protect DNA from damage (67)), the newly chromatinized genetic regulatory apparatus could “relax” itself into a new state with lower intrinsic factor specificity (in this example, by an order of magnitude) without incurring any increase in GEE, while at the same time accommodating the “spike-and-slab” distribution of target gene expression levels at lower cost of selection (68).

### Combining activation with repression enhances the benefits of chromatin-based regulation

Our analysis thus far has assumed that all TFs behave as activators. We now relax this assumption and allow each gene to be regulated by both activators and repressors. Specifically, we stipulate that each gene is regulated by three factors: one multi-target factor, always an activator; one single-target activating TF; and one single-target repressing TF. Thus, if we have *M* genes, we now have 2*M* total single-target TFs. Many eukaryotic TFs possess binding domains separate from the activating or repressing domains, therefore we consider the activating or repressing behavior to be inherent to the factor, not to the site to which it is bound. The single-target activating and repressing TFs target separate sites, and nontarget binding is permitted across all TF sites independent of the nature of their targeting factor. The dissociation rates for target and nontarget binding remain the same for all factors. We adopt a simple model for repression whereby a single bound repressor completely counteracts any activation. Thus, in free DNA, a gene is expressing only if the multi-target factor site is bound, at least one activator (which could be a multi-target factor) is bound to a single-target TF site, and no site (including the multi-target factor site) is bound by a repressor (Figure 4A). In chromatin, a gene expresses only if chromatin is permissive, at least one activator is bound, and no repressors are bound.

**Fig. 4.**
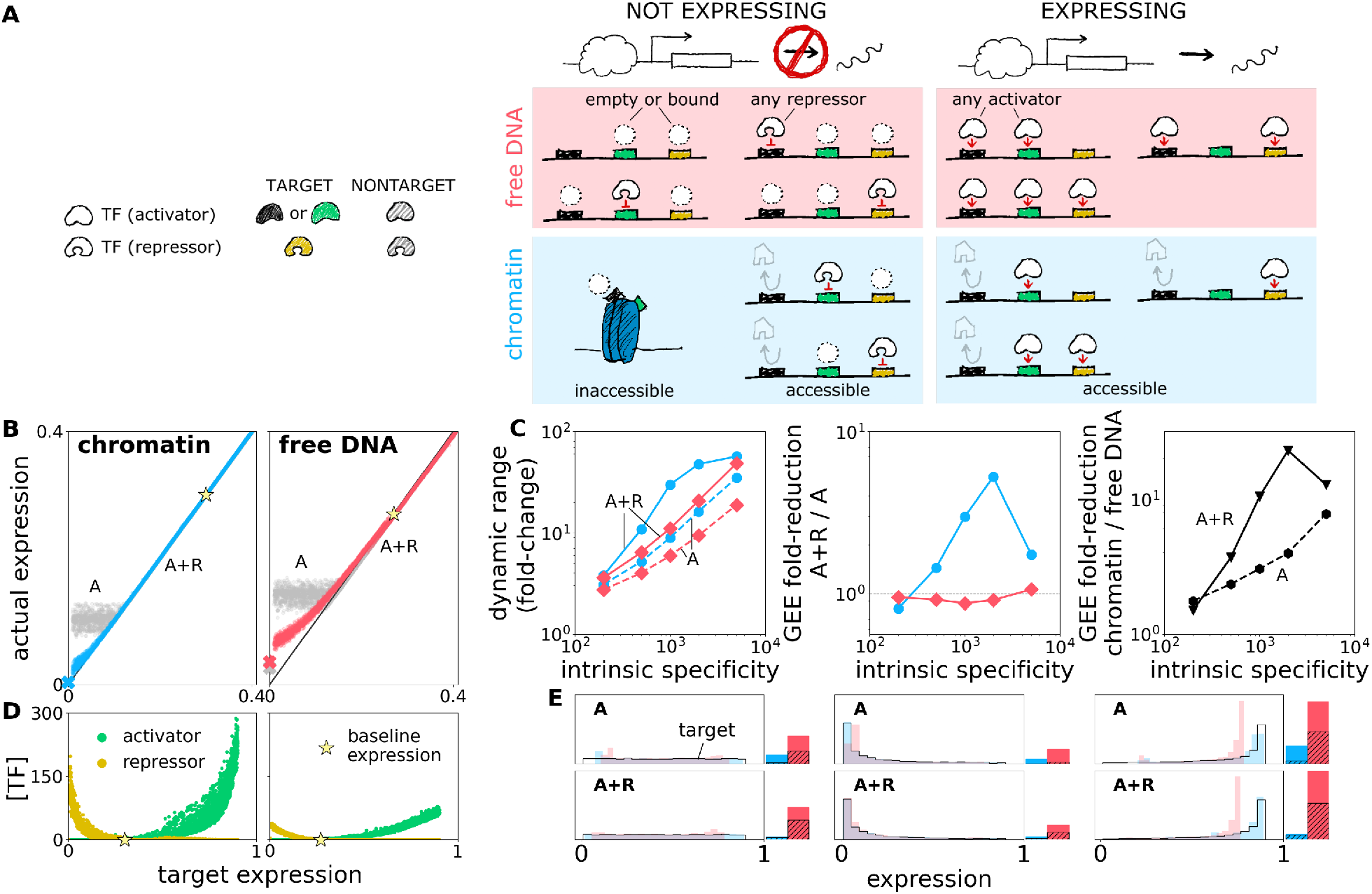
Adding repressors boosts the expressive capacity advantage of chromatin over free DNA. (**A**) A schematic of bound factor combinations that do or do not lead to gene expression. (**B**) Target vs. actual expression for simulations with activators only (as in Figure 2, “A”, gray) and with repressors (“A+R”, color) show that repressors improve accuracy at low target expression levels. (**C**) Repressors improve the expressive capacity of chromatin, but have mixed effects on free DNA. Left, the dynamic range under both scenarios is larger when repressors are present (“A+R”, solid) than when they are not (“A”, dashed), but increases more for chromatin (blue) than for free DNA (red). Middle, the fold-reduction in error when using activators and repressors vs. the activators only is typically large for chromatin (blue) but may even lead to a GEE increase for free DNA (red). Right, the fold-reduction in error upon switching to chromatin is greater when repressors are present (“A+R”, solid) than when only activators are used for regulation (“A”, dashed). (**D**) Concentrations of single-target activator (green) and single-target repressor (gold) vs. target expression level of the corresponding gene. Both concentrations reach 0 around the same target expression level, or “baseline”, where nontarget binding by activators and repressors fully accounts for expression in the corresponding genes. Expression of individual genes is modulated above this level by activators, and below it by repressors. (**E**) A regulatory architecture employing both activators and repressors (“A+R”, bottom three panels, vs. activator-only architecture “A”, top three panels) enables chromatin scenario (blue) to accommodate various target distributions of expression levels (black outlined distributions) with minimal GEE, compared to free DNA (red), whose error is significantly higher and strongly dependent on the desired gene expression level distribution. GEE with concentrations optimally adjusted to each target distribution is shown at right for each distribution, following plotting conventions of Figure 2B.

We reasoned that adding repressors to either scenario may serve to counteract the overexpression of lowly expressing genes due to nontarget binding. To ensure such an effect would be clearly visible in our simulation results, we lowered the minimum target expression level for ON genes from 0.09 to 0.009, thereby extending the target dynamic range by an order of magnitude, from 10-fold analyzed in Figure 2, to 100-fold. Simulations for this 100-fold modulation task clearly show that adding repressors indeed improves the accuracy of expression in the low-expression regime for both chromatin and free DNA (Figure 4B). Yet while adding repressors improves the dynamic range for both free DNA and chromatin, it decreases GEE only in the chromatin scenario, and actually *increases* the GEE in the free DNA scenario (Figure 4C). As suggested by previous analyses (9), this counterintuitive effect likely emerges due to the fact that adding new TFs requires new binding sites, which introduce new opportunities for nontarget interactions that limit expressive capacity. With the exception of the lowest tested intrinsic specificity (*S* = 200), chromatin compensates for this deleterious expansion of nontarget interactions by the orthogonal PF regulation and the PFs’ proofreading capacity. Thus, the opposing effects of adding repressor-mediated regulation to free DNA further widen the expressive capacity advantage for the chromatin scenario (Figure 4C).

To understand how repressors contribute to improved regulation, we plotted the optimal concentrations of single-target regulatory factors for each gene at each target expression level (Figure 4D). We observe a clear division of labor between activators and repressors: genes with high expression levels are regulated almost entirely by activators, while genes with low expression levels are regulated almost entirely by repressors. This observation implies that repressors act almost exclusively to counteract nontarget activation. The “crossover point” between regulation by repressors vs. regulation by activators—that is, the target expression level at which both repressor and activator concentration are nearly 0— indicates the baseline (“leaky”) expression level established by the combined nontarget binding activity of repressors and activators at a gene locus (Figure 4D). Subsequent modulation of specific repressor or activator concentrations therefore serves to tune the expression away from this baseline to the desired level for each target gene.

Lastly, we explored the ability of the two regulatory architectures to generate different distributions of target expression levels for ON genes. When we bias this distribution away from uniform, to favor either low or high values of gene expression, the GEE in the chromatin scenario barely changes and stays very low, while it varies substantially and is generally much higher in the free DNA scenario (Figure 4E). Thus, regulation that combines activation and repression in the chromatin scenario increases the flexibility of the regulatory apparatus: in addition to extending the target dynamic range, it can accommodate more general variations in the desired distribution of expression levels.

### Robustness to regulatory factor concentration fluctuations

In living cells, stochastic fluctuations in regulator concentrations may introduce variation in downstream gene expression levels. To examine the effect of regulator variability on expressive capacity, we start from the optimized solutions developed in the previous sections and incorporate random variation into the optimized regulator concentrations, then calculate the resulting GEE. Specifically, for each globally optimized solution that generates a target gene expression pattern, we add multiplicative noise to each optimal factor concentration *c*^*^ of regulatory factor as

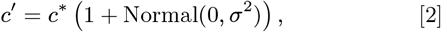

such that *c*^*′*^ is normally distributed at *c*^*^ with standard deviation *c*^*^*σ*. In this way, single- and multi-target factors are perturbed relative to their individual optima and we can vary the fluctuation level by changing the parameter *σ* (see Section S2.2 for details).

Figure 5 shows the results for fluctuations introduced to the 100-fold modulation task with a uniform distribution of target ON expression levels when both activators and repressors are present. As expected, larger fluctuations significantly decrease the accuracy of regulation and thus increase the GEE (Figure 5A, top). While chromatin still allows for smaller expression errors, it is affected more strongly by concentration fluctuations at high intrinsic specificity of regulatory factors. Surprisingly, we find that there is an intermediate level of specificity at which chromatin maximizes its advantage over free DNA (Figure 5A, bottom).

**Fig. 5.**
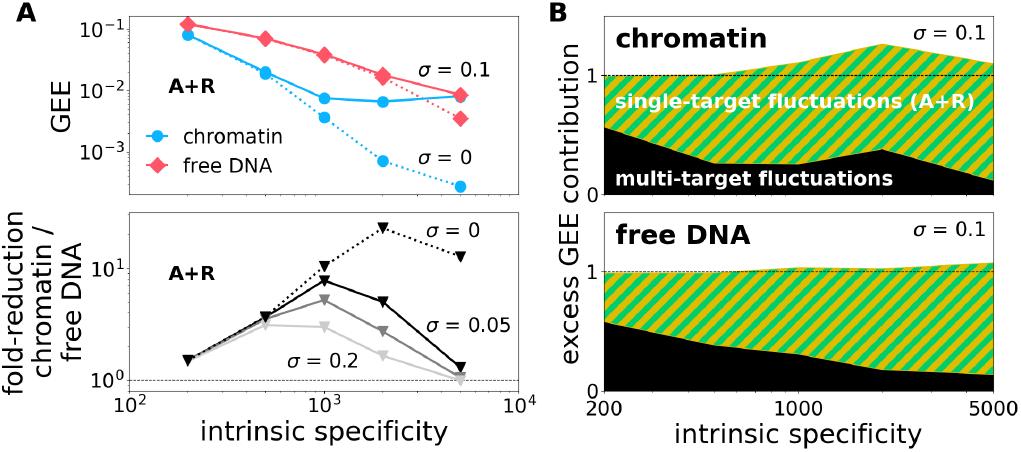
Robustness to regulatory factor concentration fluctuations. (**A**) Adding fluctuations to all single-target and multi-target factor concentrations deteriorates GEE (top) at higher levels of intrinsic specificity, more dramatically so for chromatin (blue) than for free DNA (red). Increasing fluctuation strength *σ* shifts the peak fold-reduction in GEE of chromatin vs. free DNA toward lower levels of intrinsic specificity (bottom; original noise-free results shown in dashed line). (**B**) Relative increase in GEE due to fluctuations can be attributed primarily to single-target factors. Area plots show GEE when only single-target factors (striped) or only multi-target factors (black) fluctuate with *σ* = 0.1, normalized to the GEE when all factors fluctuate simultaneously (dashed line). Results for all panels are shown for the 100-fold modulation task using activators and repressors (“A+R”), with uniform distribution of target ON expression levels.

Do multi-target or single-target factors contribute more to the degradation in accuracy upon introducing fluctuations? We added fluctuations either only to multi-target or only to single-target factors and compared the GEE between the two cases. Figure 5B shows that, across most of the specificity range, fluctuations in single-target factor concentrations result in larger expression errors than fluctuations in multi-target factor concentrations (see also Figure S4). A detailed analysis reveals the reason for this observation (Section S2.2). In optimal solutions, the multi-target factors are generally tasked with either keeping the genes entirely OFF or permitting expression. This means that the multi-target factors (and especially the PF system in chromatin scenario) operate closer to saturation, where fluctuations in the factor concentration do not propagate to gene expression. In contrast, the single-target TF system in both scenarios needs to fine-tune the expression levels of ON genes and is thereby more sensitive to fluctuations (Figure S2, S3).

Taken together, our results suggest that chromatin is well-suited when selection for extremely reliable ON/OFF control over gene expression is essential. Precise expression control is possible, but seems to require a corresponding precision in TF expression levels. As a future research direction, we note that the entire regulatory architecture and concentrations could have been optimized in the presence of fluctuations (rather than in absence, as in the setup we report here), possibly leading to an identification of noise-robust chromatin regulatory regimes.

## Discussion

Chromatin is often regarded as a central innovation in eukaryotes, but much remains to be understood concerning its implications for the global structure and strategy of gene regulatory programs. Here, we argue that chromatin, when understood as an integral and active component of gene regulatory architecture, allows for a substantially more accurate and scalable control of gene expression than is possible using TFs alone.

We motivated our study with the paradoxical observation that eukaryotes generally have much longer genomes than prokaryotes, but binding sites for transcription factors in metazoans appear to be less specific than in prokaryotes (12). This lack of specificity could lead to highly deleterious “crosstalk” in metazoan gene regulation: theory suggests that cooperative or combinatorial regulation schemes operating at equilibrium might be insufficient to ensure reliable expression control (9). Changes to effective genome size due to chromatinized sequence inaccessibility also cannot fully explain this paradox (6). Its possible resolution is a two-layer gene regulatory architecture that we propose in our work. The first layer of the architecture, namely proofreading-based modulation of chromatin accessibility, sets the discrete, ON/OFF state of the gene; the precise expression levels of ON genes are subsequently quantitatively modulated by activator and repressor TFs, acting as the second layer of the architecture. This paradigm of an “on-off switch followed by a rheostat” is well-suited to generate the “spike-and-slab” distribution of expression levels across genes (Figure 1C), with strong clustering of OFF genes across cell types and states. It is also consistent with recent experimental results showing that random stretches of DNA inserted into the genome are significantly more likely to express a reporter if a known PF binding site is included in the sequence (69).

An essential part of our approach is a global, systems-level analysis of end-to-end regulatory performance of both layers. This reveals a key insight responsible for a large boost in performance of chromatin-based regulation: Because OFF genes can be kept reliably silent via proofreading-based PF regulation, the regulatory system can indeed operate with TFs of lesser specificity and productively co-opt their non-target binding. This sets a leaky baseline expression level for all genes that targeted TF binding then up- or down-modulates to a precise desired level. Without reliable silencing of expression by PFs via chromatin, such co-opting necessarily leads to a possibly deleterious unwanted induction of OFF genes, as can be seen in our free DNA scenario.

Our qualitative conclusions concerning the benefits of chromatin are not specific to a particular molecular pathway, but apply broadly to any kinetic proofreading mechanism whereby increasing concentrations of regulatory factors favor transcriptionally permissive states. The parameter values we selected for our simulations were based on the kinetics of nucleosome repositioning in response to remodeler recruitment; however, higher levels of chromatin organization also show dose-dependent responses to regulator concentrations commensurate with our key assumptions. For example, experiments and modeling indicate that compacted chromatin can decompact with increasing concentrations of targeted activator (42, 43) (though whether compacted chromatin is truly less accessible than non-compacted chromatin is a subject of continuing debate (40, 70)). Absent from our model is explicit regulation of chromatin closing or the maintenance of transcriptionally repressive states, for which experimental evidence suggests repressor complexes such as Polycomb may be essential (36, 71–73). As in the case of TF repressors, we expect these elements to change the quantitative but not the qualitative conclusions of the study.

Perhaps the most notable omission in our analysis is that of dynamics. The chromatin regulation mechanisms discussed above differ substantially in their rates: Chromatin decompaction occurs on a timescale that is rapid compared to the length of a cell cycle (1.5 h vs. 20 h) (43), but it is still orders of magnitude slower than nucleosome repositioning (on the order of s or min). Similarly, TF binding and unbinding events as well as transcription initiation are faster than most chromatin remodeling processes (74). Therefore, in systems where speed is more important than accuracy in absolute expression levels, we may expect TFs to remain in use despite the reduced global error facilitated by chromatin-based regulation.

Because we evaluate expression levels at steady state, our results most directly apply to genes that must stably maintain a constant expression level over a long timespan relative to the rates of transcription and protein decay. The most salient biological analogs include “housekeeping” genes that carry out basal functions in all cells, as well as tissue- or cell type-specific genes that show little cell-to-cell or temporal variation in transcript levels (75–77). In addition to constant expression, three features make stably expressed genes (SEGs) particularly suitable for analysis with our paradigm. First, SEGs vary in the absolute expression levels they maintain, with one analysis revealing a 3–4-fold difference between the 25th and 75th percentiles of expression, with an absolute dynamic range of *>*50-fold overall (78). Researchers have also hypothesized means for graded control of housekeeping genes specifically, for example, through the activity of corepressors (79). Second, chromatin remodeling may silence some sets of SEGs during the development of particular cell lineages (80). Third, experimental evidence from embryonic fruit flies suggests that housekeeping genes (expressed in all cells) rely on chromatin remodelers to position nucleosomes downstream from transcriptional start sites, rather than to regulate which TFs can bind. In contrast, developmental genes rely on a different set of remodelers to maintain accessibility at enhancers (81). Thus, housekeeping genes and cell type-specific SEGs may offer a natural comparison between two alternative regulatory schemes, one of which (housekeeping) more closely resembles the free DNA scenario, and the other of which (developmental) more closely resembles the chromatin scenario.

Whether in chromatin or free DNA, an essential feature of our model is nontarget binding by regulatory factors, i.e., binding weakly to enhancers that do not regulate the factors’ central, “functional” targets. The extent to which weakly binding factors are essential, as opposed to merely consequential, for transcriptional regulation remains a subject of open discussion (10, 13, 82–84). Evidence for biologically significant weak binding emerges from studies on TF co-occupancy at enhancers, which show high prevalence in regions of accessible chromatin even between TFs with divergent binding site preferences and disparate functions (85). Weak but functional regulatory interactions have also been inferred in developmental networks, which appear to rely to a high degree on “non-canonical” TF binding interactions to establish cell type identity (86).

Our model suggests that genes may cope with unwanted weak binding by co-opting it to establish baseline expression levels for genes. Though the “leaky” expression in our model arises from uniform low-rate binding of nontarget TFs to enhancer binding sites, in practice TFs will vary along a continuum of specificity in their binding to different DNA sequences. Furthermore, it is likely that molecular steps between TF binding and transcription initiation can filter out signals emerging from binding of sufficiently low specificity (15, 87, 88) or of TFs lacking the appropriate cofactors. One prediction from these observations is that individual genes may establish baseline expression levels from subsets of the TFs that weakly bind their enhancer sequences, leading to a continuous distribution of baseline expression levels across genes. At least some data do indicate such distributions for genes expressed at less than one transcript per cell (i.e., for putatively OFF genes) (89), although we cannot rule out either natural stochasticity or binding of specific TFs in erroneously open chromatin as the cause.

How does the notion of baseline expression we propose relate to previous discussions of biologically significant weak binding? We postulate that whether a given weak interaction is “functional” may be a more nuanced question than whether large changes to TF concentration noticeably modulate a gene’s expression. In particular, some weak interactions may be *interchangeable*, in the sense that it may not matter to an enhancer *which* weakly binding TFs are present so long as *some* are. Such agnosticism to TF identity could conceivably improve the robustness of expression to uncorrelated local variations in the concentrations of weakly binding TFs.

An example of interchangeability comes from evidence that collectives of TFs can compete with nucleosomes to bind DNA and potentially activate genes (56). In particular, genomic analysis of yeast indicates that regions of DNA predicted to have high levels of nonspecific DNA binding are depleted of nucleosomes, including at the majority of promoters (90). Furthermore, DNA footprinting studies in mouse enhancers suggest simultaneous binding at regions of nucleosome competition is largely independent of TF identity and generally does not require direct TF-TF interaction (91). It would be interesting to couple such assays with measurements of the resulting gene expression driven by enhancers of interest, particularly if their strongest-binding TFs could be knocked out. Under a hypothesis of baseline expression driven by weak binding, we would expect identity-independent binding to induce some level of residual regulatory activity after knockout. A caveat to this approach is that it would not detect situations in which the activity of interchangeable weakly binding TFs is negligible or absent unless specific TFs are also present, for example, if the specific TF is an activator and the weakly binding TFs are repressors.

Nontarget binding may be co-opted or tolerated when relatively few genes are actively regulated. As more enhancers and corresponding genes become accessible, however, the unwanted effects of nontarget binding become harder to avoid. Some stem cells, for example, express genes specific to multiple possible differentiated lineages simultaneously, without performing the functions of cells belonging to these lineages (92). Such promiscuous expression might arise simply through the accessibility of regulating enhancers even if their specific binding factors are absent. Interestingly, many cancers also express embryonic stem cell markers or pluripotency factors, including PFs like OCT4 (93, 94). This may help to explain some of the phenotypic plasticity of cancerous cells as follows (95): Suppose a differentiated cell escapes into a cancerous state simultaneous with the erroneous expression of PFs. At the moment of escape, the regulatory concentrations of TFs present in the cell were optimized for the differentiated cell—that is, in an environment in which the appropriate genes were silenced. If these genes are suddenly made accessible to regulation, we might expect a higher probability of spuriously activating an unwanted transcriptional program than in an endogenous stem cell, where the regulatory concentrations would be optimized assuming these genes were accessible. Indeed, stem cells may maintain appropriate function despite susceptibility to high levels of nontarget binding due to a reliance on signaling factors secreted by other stem cells; the tumor-like behavior of cancer cells can be suppressed by these same signaling factors (96).

Chromatin may not have emerged for the purpose of regulating genes, but subsequent evolution has dramatically expanded its capacity to do so. With our work, we have aimed to provide comparative, quantitative arguments to help interpret the qualitative differences in regulatory schemes observed between prokaryotes and their eukaryotic descendants. It is our hope that further research will eventually explain, not merely the varying capacities of these genetic regulatory systems, but how their interaction with evolutionary pressures eventually gave rise to the complex, multicellular organisms that have so long excited the scientific imagination.

## Materials and Methods

Code for all simulations is available on GitHub at https://github.com/officerredshirt/network_crosstalk.

## ACKNOWLEDGMENTS

MLP was supported by the European Molecular Biology Laboratory Interdisciplinary Postdoc Programme (EIPOD4 fellowships), cofunded by Marie Sklodowska-Curie Actions (grant agreement number 847543). J.C. and M.L.P. were supported by EMBL Core Funding and Theory@EMBL. GT acknowledges the support of the Human Frontiers Science Program Grant (RGP0034/2018). The authors would like to thank the members of the Crocker and Tkačik groups, especially Natalia Misunou, Michal Hledík, and Réka Borbély, for helpful feedback and discussion. The authors also thank EMBL IT Services for the use of HPC resources.

## Notes

https://github.com/officerredshirt/network_crosstalk

### Competing Interest Statement

The authors have declared no competing interest.

