## Supplementary Information for "Chromatin enables precise and scalable gene regulation with factors of limited specificity"

June 13, 2024

### S1 Modeling

#### S1.1 Regulatory architecture

Let $\tilde{N}$ be the number of multi-target factors and $N$ the number of single-target factors with concentrations $\tilde{c}_i$, $c_j$ for the $i$th multi-target factor, $j$th single-target factor respectively. Each of the $M$ genes is regulated by one corresponding enhancer targeted by one multi-target factor and one single-target factor. In the chromatin scenario, the multi-target factor is a pioneer factor and binds only when chromatin is repressive, while the single-target factor is a TF that binds only when chromatin is permissive. Each multi-target factor regulates a fixed number of genes $M_{cluster} > 1$ such that $\tilde{N}M_{cluster} = M$. Each single-target factor regulates exactly one gene such that $N = M$. We assume all factors have the same target dissociation constant $K_T$ and nontarget dissociation constant $K_{NT}$, whether they are single- or multi-target, but that pioneer factors do not bind sites in free DNA and transcription factors do not bind sites in inaccessible chromatin. We assume our models for chromatin opening and regulatory factor binding are stochastic and ergodic, such that the system has a single stationary distribution for the probabilities of random variables

$$\tilde{L}_q(\tilde{\boldsymbol{c}}, \boldsymbol{c}) = \begin{cases} 1, & \text{multi-target factor site for enhancer } q \text{ is bound (free DNA) or open (chromatin),} \\ 0, & \text{otherwise} \end{cases} \tag{S1}$$

and

$$L_q(\tilde{\boldsymbol{c}}, \boldsymbol{c}) = \begin{cases} 1, & \text{single-target factor site for enhancer } q \text{ is bound,} \\ 0, & \text{otherwise.} \end{cases} \tag{S2}$$

We then define the expression level for gene $k$ as

$$g_k(\tilde{\boldsymbol{c}}, \boldsymbol{c}) = Pr\left[\tilde{L}_q(\tilde{\boldsymbol{c}}, \boldsymbol{c}) = 1\right] Pr\left[L_q(\tilde{\boldsymbol{c}}, \boldsymbol{c}) = 1\right]. \tag{S3}$$

Let $\mathcal{C}_i$ denote the $i$th cluster regulated by multi-target factor $i$, and let each gene $k$ belong to a single cluster. Recall that each cluster contains the same number of genes such that $|\mathcal{C}_i| = M_{cluster}$ for all $i$. To define a set of target expression patterns, we first designate all clusters $\mathcal{C}_i$ where $i = 1, 2, ..., C_{OFF}$ as OFF clusters and the remaining $\tilde{N} - C_{OFF} = C_{ON}$ clusters as ON clusters, without loss of generality since $K_T$ and $K_{NT}$ are shared by all factors. For a given target pattern, the target expression level $t_k$ of gene $k$ equals 0 if gene $k$ belongs to cluster $i$ where $i \leq C_{OFF}$, and is otherwise drawn from the uniform distribution $\mathcal{U}(D_{min}, D_{max})$ from $D_{min} > 0$ to $D_{max} < 1$.

### S1.2 Free DNA binding

We adopt a thermodynamic equilibrium model for regulatory factors binding to free DNA. Here, we report the probability that a single binding site is occupied by a target or nontarget factor, which gives us the distribution of $L_q$ for both scenarios as well as $\tilde{L}_q$ for the free DNA scenario.

Let $K_T$ be the dissociation constant for target binding and $K_{NT}$ the dissociation constant for nontarget binding. Define

$$C = \sum_i c_i \tag{S4}$$

as the sum total of all free DNA binding factor concentrations (i.e., all $\tilde{N} + N$ factors in the free DNA scenario or just the $N$ single-target factors in the chromatin scenario). Then the probability $p_{on}$ that a site targeted by factor $k$ is occupied is given by

$$p_{on} = \frac{\left(\frac{C-c_j}{K_{NT}}\right) + \left(\frac{c_j}{K_T}\right)}{1 + \left(\frac{C-c_j}{K_{NT}}\right) + \left(\frac{c_j}{K_T}\right)}, \tag{S5}$$

as derived in detail in prior works [1].

In defining the gene expression $g_k$ as the product given in Equation S3, we follow the common assumption that gene expression is directly proportional to binding site occupancy [1, 2], which has some experimental support [3]. Note that in the free DNA scenario this assumption implies that both sites must be occupied to initiate transcription and also that there are no energetic interactions, directly or indirectly, between the multi-target and single-target bound factors.

### S1.3 Chromatin opening: kinetic proofreading

Models in the literature suggest that chromatin opening follows a kinetic proofreading scheme to ensure accurate targeting of remodelers to appropriate genomic sites [4–6]. Kinetic proofreading is a nonequilibrium mechanism for enhancing the discrimination between correct and incorrect substrates in a chemical reaction, reviewed in [7]. Here, we adopt a relatively simple implementation of proofreading for chromatin opening, pictured in Figure S1. In this model, chromatin-binding factors bind and unbind chromatin with fixed on and off rates. In general, the on rate may be the same for both target and nontarget factors, but the off rate is much larger for nontarget factors. After an initial binding event, an irreversible energy-consuming step takes place that does not yet render chromatin permissive. The bound factor must then remain bound long enough for a second irreversible step to render the chromatin fully permissive. By not transitioning immediately into the permissive state upon binding, the kinetic proofreading step essentially gives the factors a second chance to dissociate, which nontarget factors will do at a much higher rate than target factors.

The transition weight matrix for the scheme pictured in Figure S1 is given by

$$Q = \begin{bmatrix} \cdot & c_S\hat{k}_S^+ & c_{NS}\hat{k}_{NS}^+ & 0 & 0 & 0 \\ \hat{k}_S^- & \cdot & 0 & k_{neq} & 0 & 0 \\ \hat{k}_{NS}^- & 0 & \cdot & 0 & k_{neq} & 0 \\ \hat{k}_S^- & 0 & 0 & \cdot & 0 & k_{neq} \\ \hat{k}_{NS}^- & 0 & 0 & 0 & \cdot & k_{neq} \\ r^- & 0 & 0 & 0 & 0 & \cdot \end{bmatrix} \tag{S6}$$

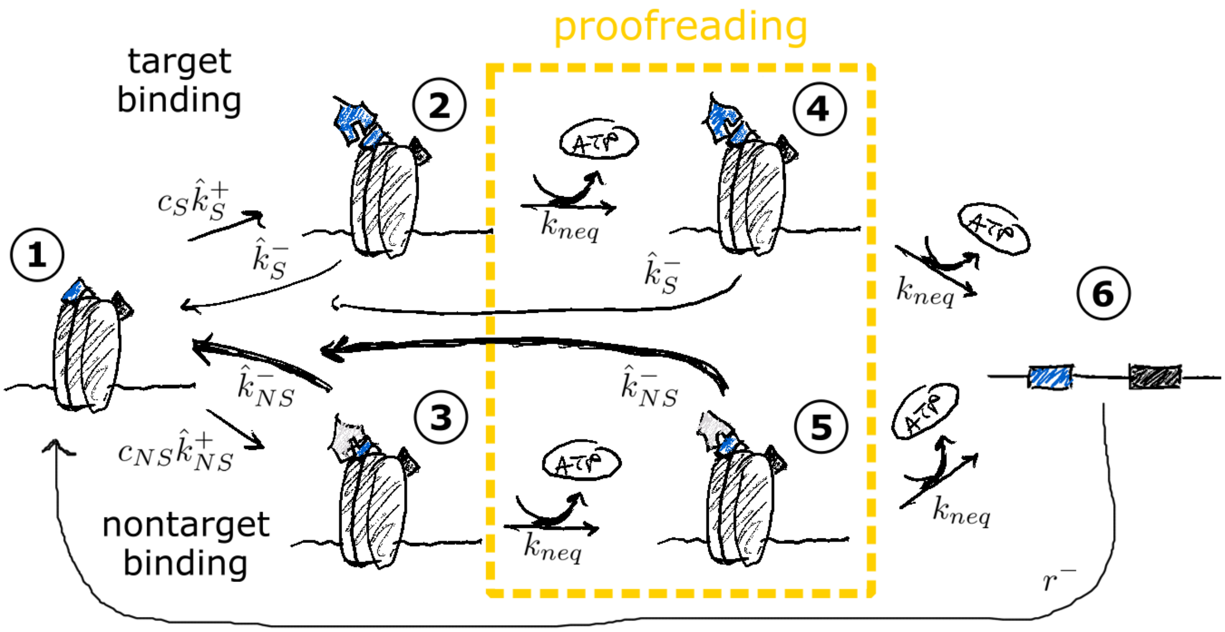


Figure S1: Kinetic proofreading model for chromatin opening with target (blue) and nontarget (gray) binding. Arrows with abutting arced arrows indicate energy-consuming steps. States are numbered to match the transition matrix in Equation S6.

where the diagonal entries are such that all rows sum to 0, and the hat designates rates associated with regulatory factors binding chromatin (vs. free DNA). The stationary solution $\boldsymbol{\pi}$ satisfies

$$\boldsymbol{\pi} = \boldsymbol{\pi} Q. \tag{S7}$$

Parameter values for simulations are given in Table S1.

### S2 Simulation

#### S2.1 Optimization to penalize nontarget binding

The error fraction for TF binding is defined as the ratio of the probability that a nontarget factor is bound to the probability that any factor is bound, which in our equilibrium model is given by

$$E_i = \frac{\frac{C-c_i}{K_{NT}}}{\frac{C-c_i}{K_{NT}} + \frac{c_i}{K_T}}. \tag{S8}$$

The error fraction for binding in chromatin is defined as the fraction of the transition rate from closed to open that is due to nontarget binding, i.e.,

$$\tilde{E}_i = \frac{k_{neq}\pi_5}{k_{neq}\pi_5 + k_{neq}\pi_4} = \frac{\pi_5}{\pi_5 + \pi_4}, \tag{S9}$$

where $\boldsymbol{\pi}$ is the stationary solution and the states are numbered as in Figure S1.

| **Parameter** | **Value** | **Source** | **Source description** |
|---|---|---|---|
| $r^-$ | 0.0002 $s^{-1}$ | [8]; [9] | (mouse) time to close on PF depletion: 30 min–2 hr; (yeast) nucleosome turnover time: $\sim$1 hr |
| $\hat{k}_S^+$ | 0.0005 $s^{-1}$ $nM^{-1}$ | [10] | specific PF binding rate (yeast) 0.0002–0.0006 $s^{-1}$ $nM^{-1}$ |
| $\hat{k}_S^-$ | 0.005 $s^{-1}$ | [10] | specific PF dissociation rate (yeast) 0.004–0.010 $s^{-1}$ $nM^{-1}$ |
| $\hat{k}_{NS}^+$ | 0.0005 $s^{-1}$ $nM^{-1}$ | [11] | (yeast) same as specific PF binding |
| $\hat{k}_{NS}^-$ | $S\hat{k}_{NS}^-\frac{\hat{k}_S^-}{\hat{k}_S^p}$ $s^{-1}$ | [12] | intrinsic specificity $S$ as in main text from nonspecific binding dissociation constant (yeast) $\sim 10^{-6}$–$10^{-3}$ M |
| $k_{neq}$ | 0.05 $s^{-1}$ | [13] | (yeast) chromatin remodeler stable residence time (including time expended for remodeling): 4–7 s |
| $K_T$ | 10 $nM^{-1}$ | [10] | specific TF dissociation rate (yeast) 0.25–0.65 $s^{-1}$ / specific TF binding rate (yeast) 0.02–0.03 $s^{-1}$ $nM^{-1}$ |
| $K_{NT}$ | $SK_T$ $nM^{-1}$ | [11], [12] | intrinsic specificity $S$ as in main text from nonspecific dissociation constant (yeast) $\sim 10^{-6}$–$10^{-3}$ M, nonspecific TF binding rate (yeast) same as specific TF binding |

Table S1: Parameter values used in simulations. Note that the dissociation constant $K_T = 10$ as approximated from measurements in [10] is on the same order of magnitude as that measured for the fruit fly transcription factor GAGA (5.2 nM, BioNumbers Accession ID 104594).

Note that we can treat the error fractions as *probabilities* that an event (binding, chromatin opening) is due to nontarget binding. Assuming the probability of nontarget binding at the cluster level is independent of the probability of nontarget binding at the individual level—with both probabilities conditioned on a gene being expressed—the total error fraction $\epsilon_i$ for gene $i$, or the fraction of expression due to nontarget binding at the cluster or individual levels, is given by

$$\epsilon_i = E_i + \tilde{E}_i - E_i\tilde{E}_i. \tag{S10}$$

When optimizing to reduce nontarget binding, we then use the objective function

$$f(\boldsymbol{c}) = \min_{\boldsymbol{c}} \sum_{i=1}^{M} \sqrt{\frac{\left(t_i - g_i(\boldsymbol{c})(1-\epsilon_i)\right)^2 + \left(g_i(\boldsymbol{c})\epsilon_i\right)^2}{M}}, \tag{S11}$$

which counts toward achieving target expression levels only that fraction of expression which occurs due to target binding at both the cluster and individual levels, and penalizes expression that arises from nontarget binding at either or both levels.

#### S2.2 Fluctuations in regulator concentration

In this analysis, we consider the effect on the GEE of adding multiplicative noise to each optimal factor concentration $c^*$ as

$$c' = c^* \left(1 + \mathrm{Normal}(0, \sigma^2)\right), \tag{S12}$$

such that $c'$ is normally distributed at $c^*$ with standard deviation $c^*\sigma$. For this analysis, we do NOT reoptimize $c^*$ in the presence of noise; rather, we begin with $c^*$ from the deterministic case, then sample the concentrations around $c^*$ following the distribution in Equation S12 and calculate the resulting GEE per sample. For the results reported in this study, we generate 10 samples of perturbed regulator concentrations per global optimization result for each value of $\sigma$. The RMSE is calculated for each sample and then averaged together to obtain the GEE with fluctuations for a single optimization result (corresponding to a single target pattern). We may choose to allow fluctuations for all regulatory factors, or for single-target or multi-target factors only.

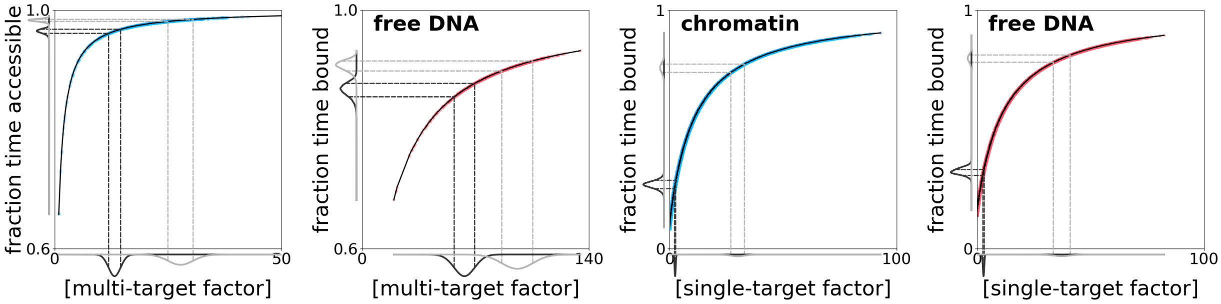


Figure S2: Modulation of accessibility in chromatin is more strongly binary than TF binding by multi-target factors, such that fluctuations about an optimal concentration produce less variation in downstream gene expression levels. Results are shown for optimal solutions (activators only) with $S = 1000$. Distributions are pictured for $\sigma = 0.1$ centered at the 20th (black) and 80th (gray) percentiles of multi-target factor concentration. Note the difference in scale on the x-axis between chromatin (blue) and free DNA (red). Chromatin's sensitivity to TF concentrations also introduces a slightly higher sensitivity to fluctuations than observed in the free DNA case.

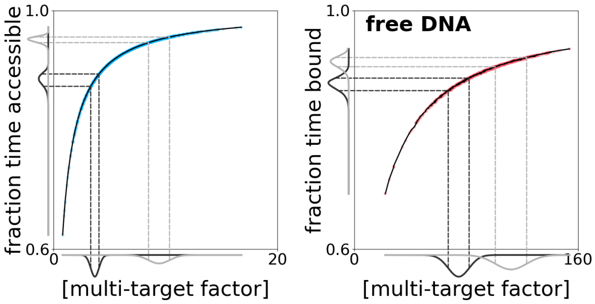


Figure S3: Same as Figure S2 but with repressors for multi-target factors only. Chromatin no longer operates as close to the saturating regime as when only activators are present, but is still less responsive than free DNA.

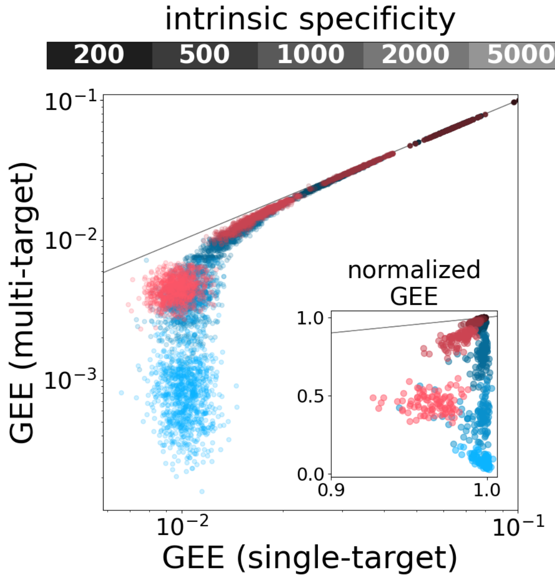


Figure S4: GEE for individual optimal patterning results (left, activators only; right, with repressors) when faced with perturbations to either single-target (x-axis) or multi-target (y-axis) factors alone. Lighter colors represent higher intrinsic specificities. Inset shows normalization to GEE when both multi-target and single-target factor concentrations are perturbed simultaneously. Gray line indicates where error contributions would be equal.

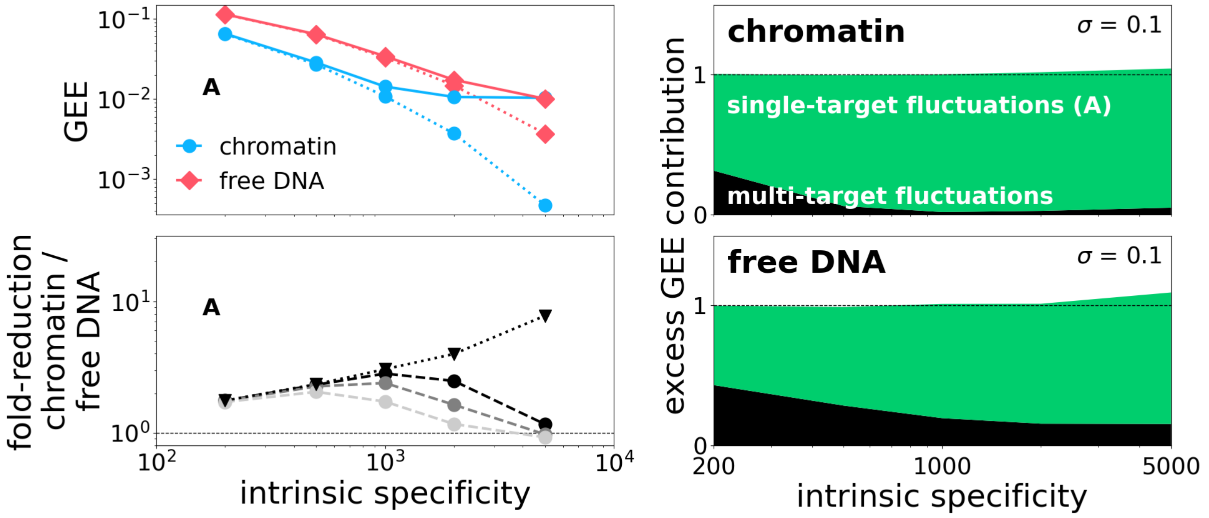


Figure S5: Same as Figure 5 but for activators only. Note that chromatin experiences less relative contribution to GEE from multi-target factors than does free DNA for all values of intrinsic specificity.

[4] R. Blossey and H. Schiessel. Kinetic proofreading of gene activation by chromatin remodeling. *HFSP Journal*, 2(3):167–170, June 2008.

[5] Ralf Blossey and Helmut Schiessel. Kinetic Proofreading in Chromatin Remodeling: The Case of ISWI/ACF. *Biophysical Journal*, 101(4):L30–L32, August 2011.

[6] Guillaume Brysbaert, Marc F Lensink, and Ralf Blossey. Regulatory motifs on ISWI chromatin remodelers: Molecular mechanisms and kinetic proofreading. *Journal of Physics: Condensed Matter*, 27(6):064108, February 2015.

[7] Hinrich Boeger. Kinetic Proofreading. *Annual Review of Biochemistry*, 91:423–447, April 2022.

[8] Michela Maresca, Teun van den Brand, Hangpeng Li, Hans Teunissen, James O. J. Davies, and Elzo de Wit. Pioneer activity distinguishes activating from non-activating pluripotency transcription factor binding sites. 2022.
